# Temporary anosmia in mice following nasal lavage with dilute detergent solution

**DOI:** 10.1101/596601

**Authors:** Thomas Gerald Mast, Kelsey Zuk, Andrew Rinke, Khaleel Quasem, Bradley Savard, Charles Brobbey, Jacob Reiss, Michael Dryden

## Abstract

Olfactory sensory deprivation induces anosmia and reduces tyrosine hydroxylase and dopamine levels in the olfactory bulb. The behavioral consequences specific to the loss of olfactory bulb dopamine are difficult to determine because sensory deprivation protocols are either confounded by side effects or leave the animal anosmic. A new method to both induce sensory deprivation and to measure the behavioral and circuit consequences is needed. We developed a novel, recoverable anosmia protocol utilizing nasal lavage with a dilute detergent solution. Detergent treatment did not damage the olfactory epithelium as measured by scanning electron microscopy, alcian blue histology, and acetylated tubulin immunohistochemistry. One treatment induced anosmia that lasted 24-48 hours. Lastly, five days of treatment reduced both olfactory bulb tyrosine hydroxylase and dopamine levels which indicates that anosmia persists between treatments. This is the first report of a sensory deprivation protocol that induces recoverable anosmia and can be paired with biochemical, histological, and behavioral investigations of olfaction.

## Introduction

The olfactory bulb (OB) is the first central structure to detect olfactory cues, providing a neural representation of the environment. Olfactory sensory neurons enter the OB glomerular layer [31] and synapse on both interneurons and output neurons [15, 32]. Thus, the glomerular circuit can modulate both sensory input and bulbar output [26]. Short axon cells within the glomerular circuit release both GABA and dopamine as neurotransmitters [19]. Dopaminergic function with the glomerular circuit is poorly defined; in vivo anesthetized studies typically find little effect of dopamine on sensory neuron activity [27]; however, short-axon cells actively regulate dopamine production to match sensory input.

Short-axon cells are particularly sensitive to sensory deprivation as measured by OB dopamine concentrations [3] and by the number of tyrosine hydroxylase (TH) interneurons [17, 18, 42]. Although olfactory sensory deprivation was originally developed as model to study cell death during ‘critical periods’ in development [5], histological and biochemical effects are similar in the adult mouse OB [24, 42]. In vitro, short axon cells can produce presynaptic inhibition [11] and center-surround inhibition [2]. Thus, reduced short axon cell activity following sensory deprivation is hypothesized to maintain ‘set-point’ of neuronal activity in a process termed homeostatic plasticity [7].

The behavioral consequences specific to OB homeostatic plasticity are difficult to determine because the methods that induce OB plasticity either destroy sensory neurons [3] or block nasal airflow and odorant access to olfactory sensory neurons [24]. Both techniques render the animal anosmic. Even when sensory neuron activity has returned, such as after removal of the nasal blockage in naris occlusion [18, 41], a slew of compensatory responses outside the OB make interpretation of behavioral and physiological data difficult [7, 16]. Therefore, a protocol is required that induces olfactory homeostatic plasticity while accommodating biochemical, histological and behavioral techniques.

Nasal lavage with a small volume of dilute detergent does not damage the mouse olfactory epithelium. Single treatments induce anosmia for approximately 24 hours with no sign of olfactory epithelium damage. Repeated treatments prolong the anosmia and reduce OB dopamine synthesis. This protocol will allow for the investigation of the olfaction during homeostatic plasticity.

## Materials and methods

### Animals

Mice were bred from breeder-pairs purchased from The Jackson Laboratories (C57BL/6J, strain # 000664). Mice were housed in a temperature and humidity controlled vivarium, on a reversed 12 hour light/dark cycle, in polycarbonate cages, with corn cobb bedding (Bed o’ cobs, The Andersons), microfilter lids with food (Purina 5015) and tap water *ad libitum*. 75 male and female mice were included across all experiments (see Figure 1 for design and below). Scanning electron microscopy included 10 mice: PBS, Triton D2 (N = 5 per group). Alcian blue histology included 6 mice: PBS and Triton D2 (N = 3 per group). Acetylated-tubulin immunohistochemistry included 6 mice: PBS and Triton D2 (N = 3 per group). The buried food task included 25 mice: PBS (N = 8), Triton D2 (N = 9), and Triton D3 (N = 8). The Habituation-discrimination task included 18 mice: PBS, D2-PBS, and Triton D6 (N = 6 per group). Tyrosine hydroxylase (TH) immunohistochemistry included 12 mice: PBS and Triton D2 (N = 6 per group). The Eastern Michigan University Animal Care and Use Committee approved the procedures (protocol #2017-079).

**Figure 1.**
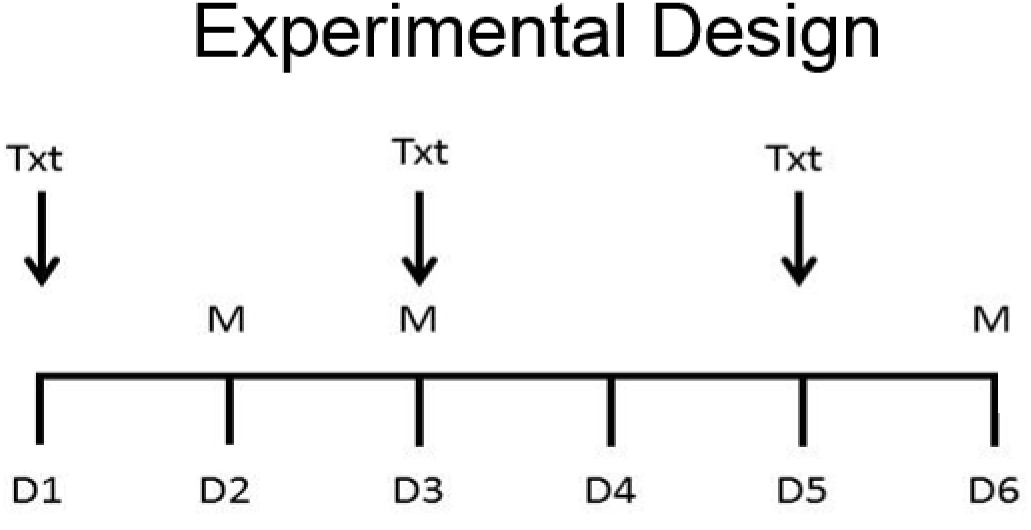
Experimental Design. Nasal lavage (TXT) with either PBS or 0.1% Triton (10 µl) began on day 1 (D1). Measurements (M) occur either 24 hrs (D2 or D6) or 48 hrs (D3) after the last treatment. Acute experiments have one treatment (D2 or D3). Chronic experiments have three treatments (D6).

### Nasal Lavage and Treatment Groups

Mice were hand-restrained, held upright, and given one, 10 microliter nasal flush of phosphate-buffered saline (PBS) or 0.1% Triton/PBS (Triton) per nare. For a negative control, some mice were hand-restrained and had a pipette placed against the nare, but were not irrigated (Scruff). Lavages were administered on days 1, 3, and 5. On days 2, 3 and 6 (i.e. Triton D6) mice were further tested for behavior, immunohistochemistry, or biochemistry (Figure 1).

### Buried Food Task

Mice were first exposed to the food (malted milk ball, Hershey’s; Hershey, PA) for two days in their home cage to prevent neophobia. Mice were food restricted for 12 hours before the task began to control for motivation and to ensure mice utilize olfaction during chamber investigation [21, 43]. To begin the task, mice were habituated to the testing chamber (a clean cage similar to the home cage but without food or water) for 10 minutes in dim red light. Mice were then transferred to a testing chamber with a buried round chocolate candy The food was placed approximately 1 cm below the bedding surface and 1 cm above the chamber bottom. The food was placed in one of the two back chamber corners approximately 2 cm from each wall. Each mouse performed two buried food trials with the corner side alternated systematically (i.e. back left then back right). Each mouse performed a third trial with the candy on top of the bedding. All mice found the candy on top of the bedding. Each trial lasted until the food was uncovered or directly investigate or after 300s. Mice were placed in the habituation chamber for 300s between trials. Time to find the buried food (two values per mouse) and success rate was measured.

### Olfactory Habituation-Discrimination Behavior

Mice were first habituated to the testing chamber (a clean cage similar to the home cage but without food or water) for 10 minutes in dim red light. Odorants (10 µl) were applied to the tip of a cotton swab and inserted 2-3 cm into the corner of the cage through the wire lid. The total time of investigatory behaviors–such as rearing and active sniffing–within 2-3 cm of the cotton swab was recorded for each one-minute presentation. The task included four habituation odor presentations (trials) to a weak stimulus (light mineral oil; Sigma), and a final, strong odor test trial (acetophenone diluted 1:1000 light mineral oil; Sigma). The inter-trial interval was three minutes.

### Anatomical Analysis and Light Microscopy

Mice, tissue, and analysis were collected and completed as previously described [23, 24]. Briefly, mice were overdosed with urethane (0.25 g/ml) perfused (PBS and 4% paraformaldehyde/PBS), then OBs were removed, post-fixed overnight at 4°C, cryoprotected (30% sucrose /PBS). To section the olfactory epithelium, skulls were decalcified in 4°C 0.5M EDTA as previously described [23]. Tissue was sectioned at 16µm with a cryostat (Minotome). OB sections were incubated with TH antiserum (1:2000; Immunostar) and olfactory epithelia with acetylated-tubulin antiserum (1:200; Sigma) followed by donkey anti-mouse 488 secondary antisera (1:400; Jackson Immunoresearch), and diamidino-phenyindole (DAPI, Fisher) in PBS (1:20,000).

Sections were imaged on a Nikon E400 equipped with a CoolSnap Myo (Photometrics; Mager Scientific) and NIS Elements. TH-immunopositive (TH+) neurons from 3 sections per OB were summed (cell counter plugin, ImageJ) from dorsal, ventral, medial, and lateral fields of view (12 total; 74,400 µm^2^ each). Acetylated tubulin immunolabeling was quantified using a corrected fluorescence method [25]. Briefly, in ImageJ, 2 regions of interest (ROIs) were selected from the cilia layer and 3 ROIs were selected from background. Two calculations were performed per slide, 1 from the septum and 1 from a dorsal recess. Two slides were quantified per mouse. Image brightness and contrast were adjusted for clarity.

### Scanning Electron Microscopy

Epithelia were prepped as before [38]. Briefly, tissue was dehydrated in a graded series of ethanol from 50%, 70%, 90%, to 100% successively for 20 minutes in each concentration. Dehydration was completed with two 20 minute washes of hexamethyldisilazane (HMDS) and an overnight period to off-gas. Tissue was mounted to a graphite tape covered metal stub and sputter coated in gold with an SPI module sputter coater 12151 (Structure Probe, Inc. West Chester, PA 19381). Tongues were examined using an Amray 1820 Digital Image Scanning Electron Microscope (AMRAY Bedford, MA) with an acceleration voltage of 5kV. Micrographs were captured as tiffs (SEMtech) and stored on a computer for analysis.

### Dopamine Detection by HPLC

Following decapitation, OBs were rapidly dissected onto dry ice, and stored at -80°C until shipment and HPLC detection. Dopamine was measured by HPLC by the Vanderbilt Neurochemistry core.

### Hypothesis Testing and Statistics

For the buried food task: Triton D2 mice were predicted to find fewer buried whoppers and to have a longer latency as compared to PBS or Triton D3. For acetylated tubulin immunohistochemistry: no difference in the corrected fluorescence was predicted. For the habituation-discrimination task: all mice were predicted to habituate between the first and fourth mineral oil trials (i.e. MO1>MO4). Triton D6 mice were predicted to fail to discriminate between the fourth mineral oil trial and the test odor trial (i.e. MO4<TEST). For TH+ cell counts and OB dopamine concentration: Triton D6 were predicted to have fewer TH+ neurons and lower OB dopamine concentration (i.e. PBS>Triton D6). Buried food trial outcome were tested with a chi-square. Latency to buried food and the corrected fluorescence were tested with one-way ANOVA. Habituation and discrimination were tested with paired one-tailed t-tests, and the alpha was set to a Bonferroni corrected p=0.025. TH+ soma and dopamine concentrations were tested with unpaired one-tailed t-tests; the alpha was set to p=0.05 (histochemical), or a Bonferroni corrected p=0.025 (biochemical). Effect sizes were calculated following Cohen (d) [6].

## Results

Anecdotal evidence suggests that small volumes of 0.1% triton damages the olfactory epithelium [36], particularly olfactory sensory neuron cilia. Therefore, we first used scanning electron microscopy to image nasal cavity 24 hours after treatment (PBS and Triton D2; Figure 2). Nasal lavage with 0.1% triton had no discernable effect on the olfactory epithelium at low magnification (compare Figures 2A and 2B) and high magnification (compare Figures 2C and 2D). As seen in the dorsal recess, nasal lavage with 0.1% triton may reduce the thickness of the overlying mucus (compare Figures 2E and 2F). However, this was not associated with obvious damage to olfactory sensory neuron cilia (Figures 2G-2L). Cilia are readily apparent following lavage with PBS at low magnification (Figure 2G) and high magnification (Figures 2I and J). Cilia are also readily apparent following lavage with 0.1% triton at low magnification (Figure 2H) and high magnification (Figures 2K and L).

**Figure 2.**
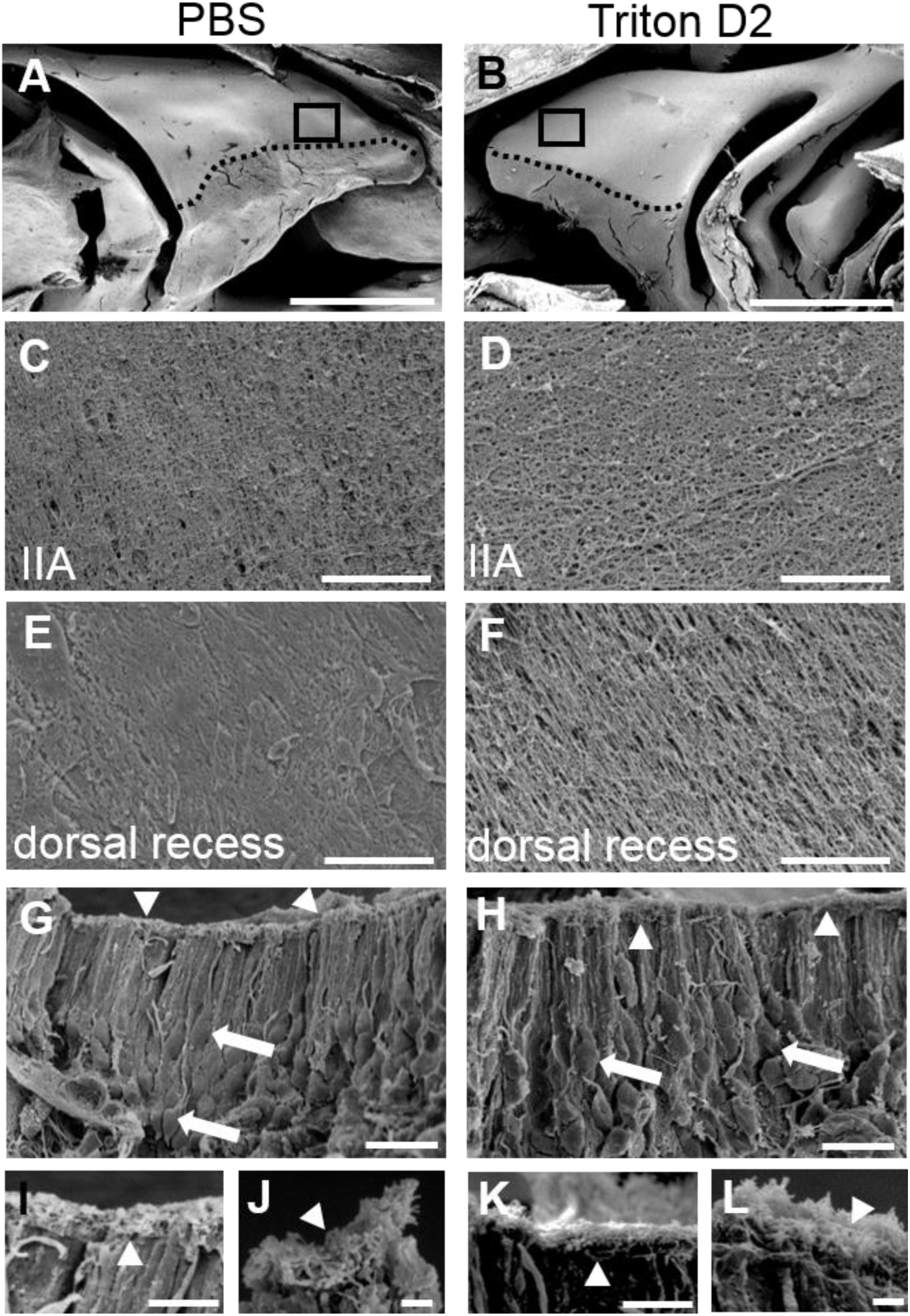
Nasal lavage with 0.1% triton does not damage the olfactory epithelium, but may wash away mucus. Scanning electron micrographs taken 24 hours after nasal lavage with either PBS (left, PBS; N=5) or 0.1% Triton (right; Triton D2; N=5). (A-B) Low magnification views (scale bars = 1mm) of the medial surface of ethmoturbinates. The respiratory epithelium on turbinate IIA is marked with a dotted line. Boxes are the fields of view in C & D. (C-D) High magnification of views (scale bars = 10 µm) of turbinate IIA. Little damage can be seen in A-D. (E-F) High magnification views (scale bars = 10 µm) of the dorsal recess. Note the thick mucus seen in E, is reduced in F. (G-H) Side view of olfactory epithelium (scale bars = 10 µm) containing olfactory sensory neurons (arrows) and covered with cilia (arrowheads). (I-L) High magnification views of olfactory cilia (I & K, scale bars = 5 µm; J & L, scale bars = 1 µm). Note that cilia are plentiful in both treatments.

Potential damage to olfactory mucus and olfactory cilia was further investigated with light microscopy. First, olfactory epithelia were stained with alcian blue to visualize mucus [14]. Nasal lavage with 0.1% triton had no discernable effect olfactory epithelium at low magnification (compare Figures 3A and 3B). At high magnification the dorsal recess (compare Figures 3C and 3D) and septum (compare Figures 2E and 2F) have comparable staining. Olfactory cilia are difficult to measure in situ [13], therefore, we used acetylated tubulin immunohistochemistry to visualize and quantify olfactory cilia [39]. Nasal lavage with 0.1% triton has no discernable effect olfactory epithelium at low magnification (compare Figures 3G and 3H). Lastly, the density of acetylated tubulin did not differ between treatments or areas (Figure 3I; F_(3,20)_ = 0.433, P= 0.731).

**Figure 3.**
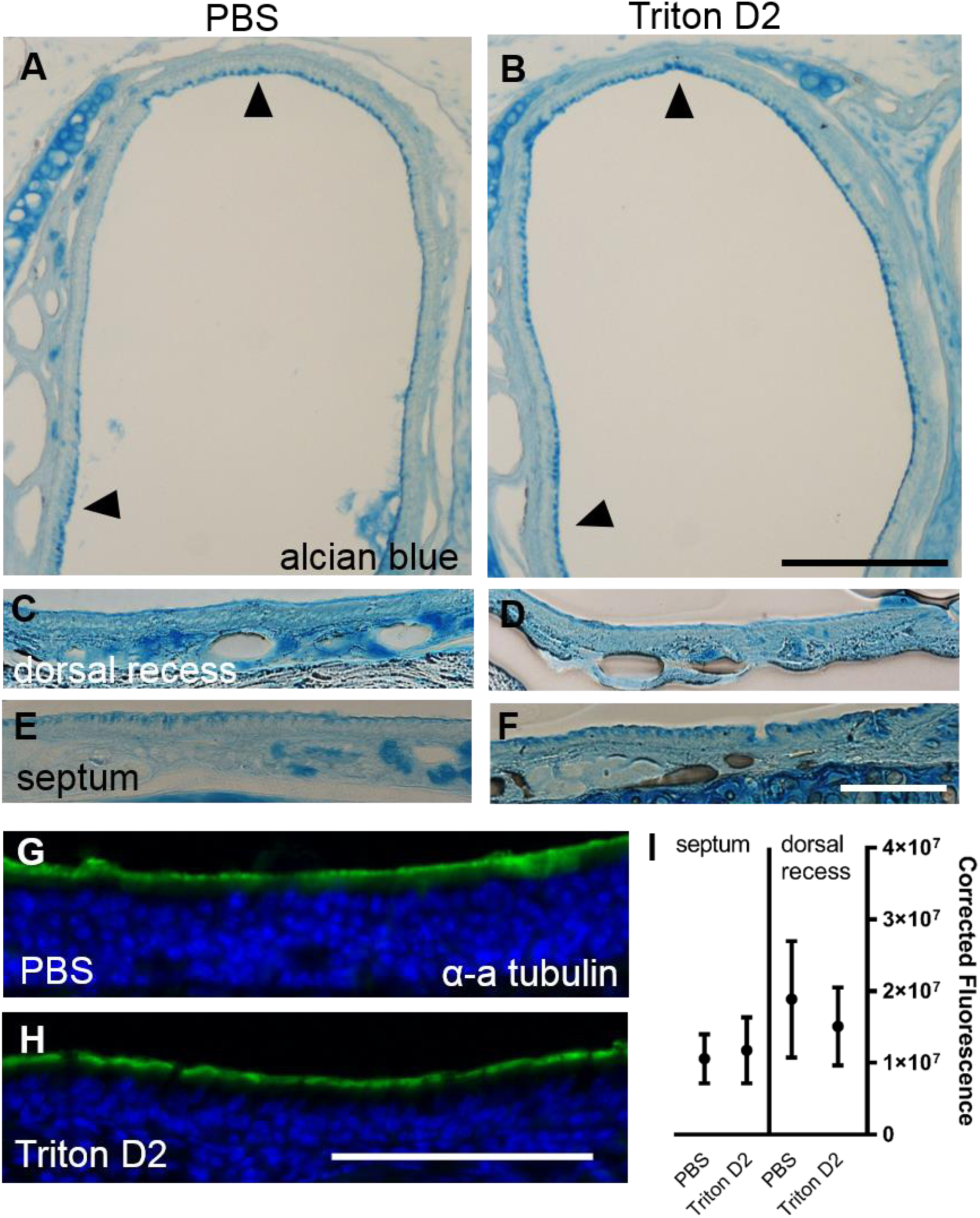
Nasal lavage with 0.1% triton does not damage the olfactory cilia. Micrographs taken 24 hours after nasal lavage with either PBS (left, PBS; N=3) or 0.1% Triton (right; Triton D2; N=3). (A-F) Transverse sections of the nasal cavity stained with alcian blue to visualize mucus (A & B, scale bars = 200 µm; C-F, scale bars = 100 µm). Dense staining is apparent on both the dorsal recess and septum (black arrowheads). In higher magnification views (C-F), little difference can be seen between treatment groups. (G-H) Acetylated tubulin immunohistochemistry (α-a tubulin; green) to visualize cilia tips in the olfactory epithelium (nuclei stained blue with DAPI) (scale bar = 50 µm). (I) Treatment with triton did not reduce α-a tubulin expression (p=0.731; N=3). Bonferroni correction used on all p-values.

Anecdotal evidence suggests that small volumes of 0.1% triton damages reduces olfactory ability for 24-48 hours [36], therefore we sought to confirm and extend this finding. First, olfactory ability was screened with the buried food task REF. Nasal lavage with 0.1% triton noticeably reduced olfactory ability (Figure 4). Triton D2 mice took significantly longer to find the buried food than the PBS-treated mice (265 versus 152 seconds; F_(2,47)_ = 13.89, p = 0.002, d = 1.28) and had fewer successful trials (8/18 versus 13/16; χ(2) = 14.23, p = 0.008). Interestingly, Triton D3 mice, which were given an extra day of recovery, had similar latency (146 versus 152 seconds; p > 0.05) and successful trials (16/16 versus 13/16) compared to PBS animals. Thus, a singular nasal lavage with 0.1% triton produces temporary anosmia.

**Figure 4.**
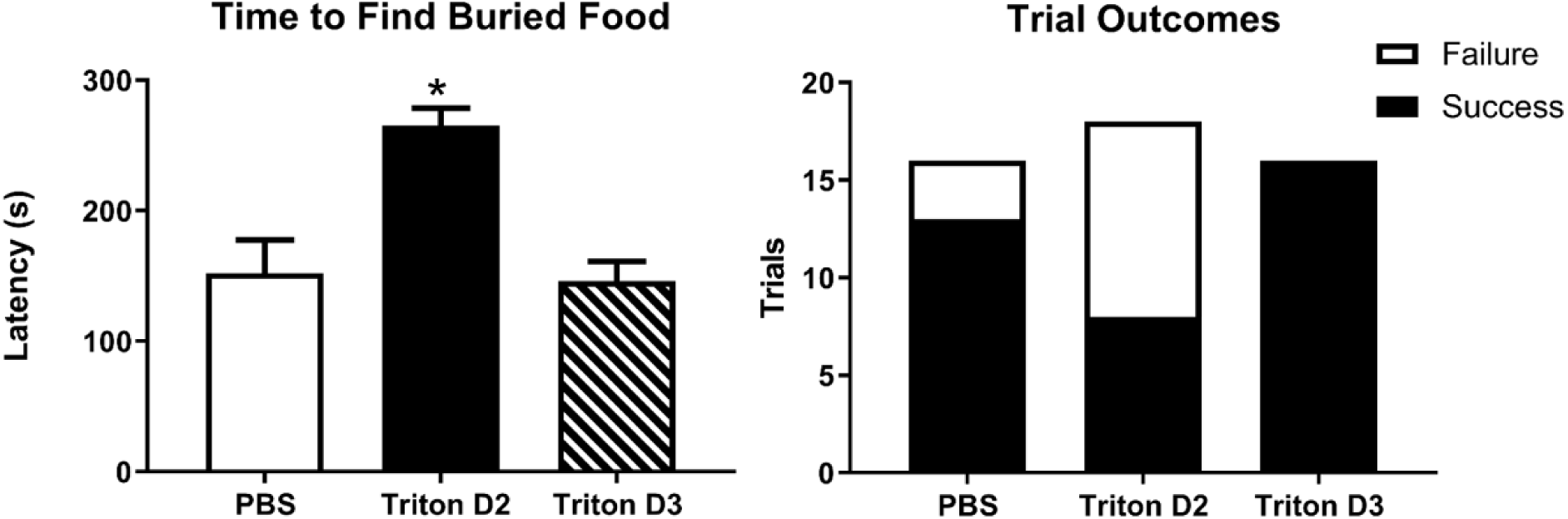
Nasal lavage with 0.1% triton induces temporary anosmia. Naive animals searched for hidden food in cage bedding (animals were used once). Speed (latency) and accuracy (success vs failure) were measured. Speed (Left) was reduced at 24 hrs (Triton D2; * p = 0.0002; Bonferroni corrected) but not at 48 hrs (Triton D3). Accuracy (Right) was also reduced at 24 hrs (Triton D2: * = p<0.0008), but not 48 hrs.

Next we tested if nasal lavage with 0.1% triton could produce homeostatic changes within the OB consistent with anosmia. Additionally, we also wanted to confirm a longer-term recovery following lavage. To test this, olfactory habituation-discrimination behaviors, and OB dopamine levels were investigated in Scruff mice that received no lavage, D2-PBS mice that received one triton lavage followed by two PBS lavages, and Triton D6 mice that received three 0.1% triton nasal lavages (Figure 5). All mice habituated to mineral oil (Scruff; t = 4.76, p = 0.0025, d = 2.87; D2-PBS; t = 4.19, p = 0.0043, d = 2.53; Triton D6; t = 4.57, p = 0.003, d = 2.76). Scruff (t = 5.15, p = 0.0018, d = 3.11) and D2-PBS (t = 3.13, p = 0.013, d = 1.88) mice discriminated between mineral oil and acetophenone (1:1000 dilution; TEST). In contrast, Triton D6 mice did not discriminate (t = 0.31, p = 0.39, d = 0.19). Scruff and PBS mice had a similar OB dopamine levels (t = 1.85, p = 0.046, d = 1.07), while Triton D6 mice had less (t = 4.56, p = 0.0005, d = 2.64). Thus, repeated triton treatment, but not a one-time triton treatment, induced chronic anosmia (i.e. failure to discriminate) and reduced dopamine production.

**Figure 5.**
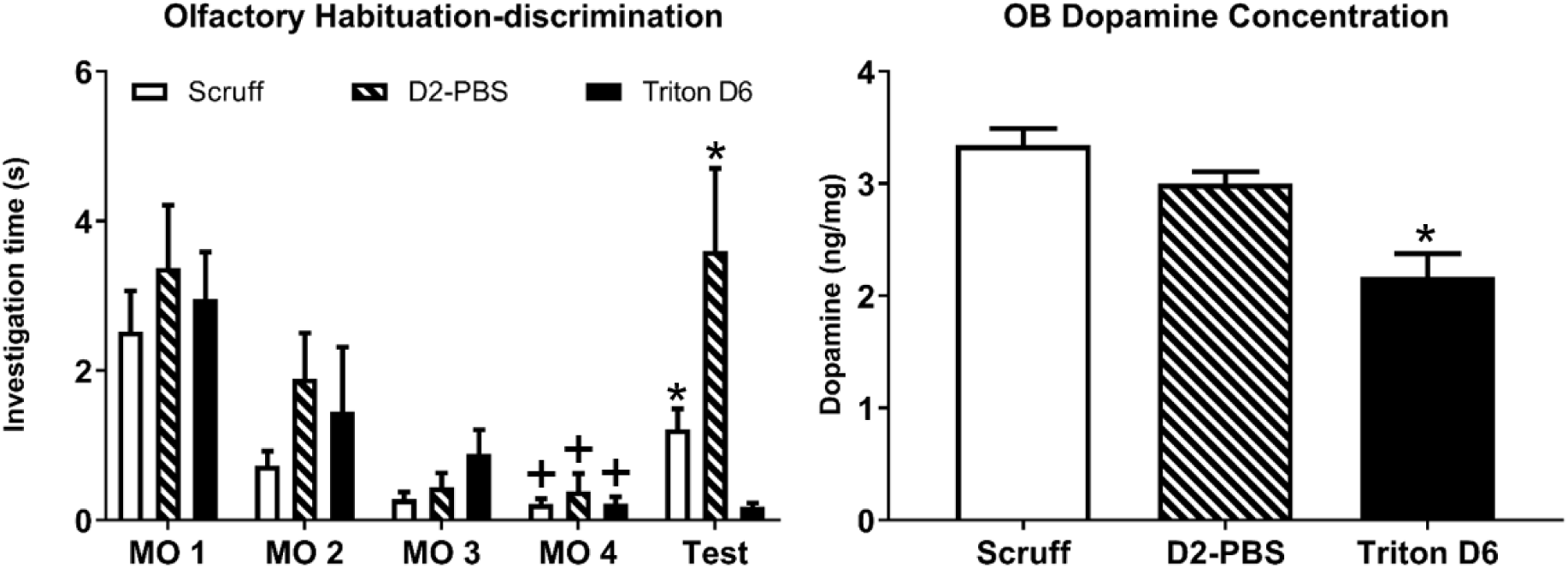
Nasal lavage with 0.1% triton induces temporary anosmia that persists with repeated treatments. Animals were presented odors on cotton swabs and investigation time was measured. (Left) All mice habituated between the first and the last mineral oil trials (MO1 v MO4; + = p<0.05; N=6). Mice treated with no nasal lavage (Scruff; white) and mice treated D2 triton and then PBS (D2-PBS; striped) discriminated between mineral oil and 1:1000 acetophenone (MO4 v TEST; * = p<0.05; N=6). Mice treated with triton three times (Triton D6 mice; black) did not. (Right) Scruff mice and mice that recovered from a singular triton lavage had similar OB dopamine content (D2-PBS vs Scruff; p>0.05). Repeated triton treatment reduced OB dopamine content (Triton D6 vs Scruff; *=p<0.05; N=6). Bonferroni correction used on all p-values.

Lastly, to verify that repeated nasal lavage with 0.1% triton reduces OB dopamine synthesis, TH expression was investigated in the OB. In a follow-up experiment, TH-immunepositive neurons were visualized and quantified in PBS and Triton D6 OBs (Figure 6). TH expression was evident in the glomerular layer (Figure 6A & 6B). Triton D6 mice had significantly fewer TH+ soma compared to PBS mice (Figure 6D; t = 3.19, p = 0.0048, d = 1.84).

**Figure 6.**
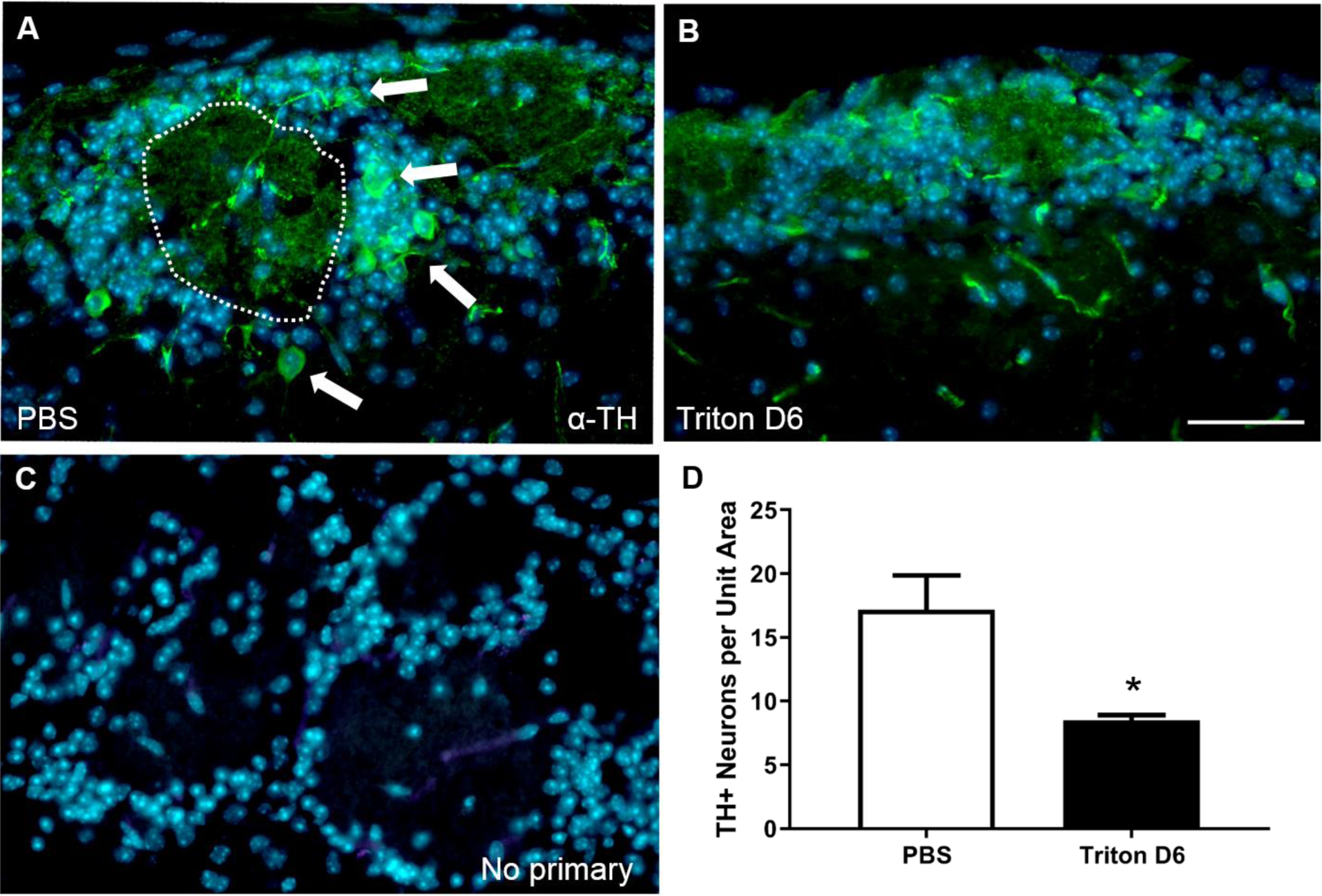
Nasal lavage with 0.1% triton reduces OB tyrosine hydroxylase (TH). TH-immunopositivity (green) was visualized in the OB glomerular layer. (A) TH+ soma (arrows) surround a glomerulus (dashed line) in a PBS-treated OB. (B) Few TH+ soma are visible in a Triton-D6 OB. Scale bar = 50 µm. (C) No primary control. Nuclei are shown for reference (blue; DAPI). (D) Bar plot: TH+ soma are significantly reduced (* = p<0.05; N=6; t= 3.2) per unit area (74,400 µm^2^) in D6 OBs.

## Discussion

We developed a protocol that combined repeated nasal flushes of a dilute detergent solution (0.1% triton) with dopamine detection, TH immunohistochemistry, scanning electron microscopy, and importantly, olfactory behavioral testing. The repeated nasal flushes induced a recoverable anosmia. These results were demonstrated in three ways. First, mice given an extra day of recovery (Triton D3) were able to find buried food—unlike mice that were tested one day following treatment (Triton D2). Second, repeated treatments with 0.1% triton reduced TH+ soma within the OB glomerular layer, consistent with reduced olfactory input [24]. Third, mice treated with 0.1% triton had reduced OB dopamine levels, again consistent with reduced olfactory input [3]. These, are the first data to measure olfactory behaviors following a loss of sensory input without blocking nasal airflow or killing olfactory sensory neurons.

Triton is known to reduce olfactory sensory neuron activity. Typically, high triton concentrations (>0.5%) are used ablate OSNs [8, 12], while dilute triton concentrations (<0.025%) are thought to spare neurons, but damage olfactory sensory neuron cilia [1]. Washing the frog olfactory epithelium with dilute triton reduces the electroolfactogram for 2-4 days [1]. Anecdotally, in mice, nasal lavage with 0.1% triton reduces the electroolfactogram for 24-48 hours [36]. Here, we confirm that 0.1% triton induces a recoverable loss of olfactory sensory neuron activity. One day after treatment, olfactory-dependent foraging is lost (i.e. Triton D2) indicating anosmia [43]. Food-finding returns the next day (Triton D3), indicating recovery from a temporary anosmia. Ciliogenesis takes about two weeks [13], and we found no evidence for cilia damage after nasal lavage in mice, therefore mouse cilia are not directly damaged the dilute triton.

The small volume of dilute triton did not damage the olfactory epithelium but may have altered the olfactory mucus. We found no evidence of epithelial damage using both light and scanning electron microscopy. Intact epithelium and cilia were evident in both techniques. While olfactory activity is dependent upon signal transduction in cilia— which are difficult to measure with SEM [13]. Therefore, cilia were visualized by immunolabeling microtubules with anti-acetylated tubulin [Williams et al., 2017] and found no difference. However, the mucus layer covering the dorsal recess did appear thinner in the triton-treated animals. Mucus is critical for olfactory ability [29]. Nasal lavage with 0.1% may have solubilized or diluted a critical mucal component such as a protein [14], sodium [37], or glucose [35]. More studies need to conducted to confirm the mechanism.

Behavioral olfactory habituation is a simple form of memory that relies on both cortical and bulbar circuits [40]. The protocol used here (1 minute odor stimulation; 3 minute inter-trial interval) is dependent on OB circuits [28]. Pharmacological studies in rodents have demonstrated that OB-dependent habituation is mediated by several neurotransmitter systems, including glutamate activation of NMDA [28], estradiol activation of ERβ [10], and norepinephrine [22]. Similar results have been demonstrated in drosophila [9]. Our results demonstrate that OB-dependent habituation fully recovers following a singular nasal lavage. Thus, future experiments that add recovery from triton-induced anosmia can investigate OB circuits during homeostatic plasticity.

OB short-axon cells form an interconnected and complex circuit within the glomerular layer where they co-release dopamine and GABA [20] and participate in center-surround inhibition [2] and gain control [34]. Sensory deprivation reduces both GABA and dopamine synthesis within the OB [30]. This is hypothesized to be a homeostatic mechanism [7] to increase sensory input (i.e. gain control). Further, reduced short-axon cell activity also decreases lateral inhibition and is hypothesized to impair OB discrimination of similar inputs [4]. Further experiments are needed to clarify the role of short-axon cells in OB odor processing and specifically need to address the potentially independent roles of GABA and dopamine.

## Conclusion

Nasal lavage with a dilute triton solution produces temporary anosmia, as indicated by a loss of olfactory-dependent food finding, olfactory discrimination, reduced TH expression, and reduced dopamine production. The protocol described here will be useful for both investigating the still undefined role of dopamine within the OB, specifically [26]; and the mechanisms and consequences of homeostatic plasticity, broadly [33].

## Acknowledgments

This work was funded by the Biology Department, College of Arts and Sciences, Honor’s College, and Provost Office at Eastern Michigan University. We thank Mr. Larkin Pence and Jim Cade for scanning electron microscope maintenance. Zach Harkey, Taylor Chick, Zach Whiddon, and Catherine Kaminski provided helpful labor.

